# A demogenetic individual based model for the evolution of traits and genome architecture under sexual selection

**DOI:** 10.1101/2020.04.01.014514

**Authors:** Louise Chevalier, François De Coligny, Jacques Labonne

## Abstract

Sexual selection has long been known to favor the evolution of mating behaviors such as mate preference and competitiveness, and to affect their genetic architecture, for instance by favoring genetic correlation between some traits. Reciprocally, genetic architecture can affect the expression and the evolvability of traits and mating behaviors. But sexual selection is highly context-dependent, making interactions between individuals a central process in evolution, governing the transmission of genotypes to the next generation. This loop between the genetic structure conditioning the expression and evolution of traits and behaviour, and the feedback of this phenotypic evolution on the architecture of the genome in the dynamic context of sexual selection, has yet to be thoroughly investigated. We argue that demogenetic agent-based models (DG-ABM) are especially suited to tackle such a challenge because they allow explicit modelling of both the genetic architecture of traits and the behavioural interactions in a dynamic population context. We here present a DG-ABM able to simultaneously track individual variation in traits (such as gametic investment, preference, competitiveness), fitness and genetic architecture throughout evolution. Using two simulation experiments, we compare various mating systems and show that behavioral interactions during mating triggered some complex feedback in our model, between fitness, population demography, and genetic architecture, placing interactions between individuals at the core of evolution through sexual selection. DG-ABMs can, therefore, relate to theoretical patterns expected at the population level from simpler analytical models in evolutionary biology, and at the same time provide a more comprehensive framework regarding individual trait and behaviour variation, that is usually envisioned separately from genome architecture in behavioural ecology.

## Introduction

Sexual selection has long been recognized as an evolutionary force shaping mating behaviours and morphological traits in populations (Darwin, 1872; Fisher, 1915; Jones and Ratterman, 2009), and has been invoked as a driving force behind speciation (Lande, 1981; Ritchie, 2007). Evolutionary biology, therefore, predicts change in the genotypic and phenotypic composition of a population due to sexual selection (Lande, 1981; Lande and Arnold, 1985; Pomiankowski and Iwasa, 1998; Tazzyman and Iwasa, 2010). Such predictions, based on population genetics, quantitative genetics, or adaptive dynamics rely on simplifying assumptions concerning mating processes and do not explicitly represent the pairing dynamics during mating.

And yet, sexual selection is fundamentally the result of complex processes involving behavioural interactions between individuals, with tremendous impacts on evolution (Bailey and Moore, 2012; Kokko, Booksmythe, et al., 2015; Moore et al., 1997; Muniz and Machado, 2018; Wolf et al., 1999). This social part of the environment is indeed one of the most dynamic sources of variation an organism might experience during its lifetime (Kent et al., 2008; Krupp et al., 2008; West-Eberhard, 1983). As a response, behavioral ecology particularly aims at understanding the effect of this variable social environment (e.g., the availability of partners of various qualities) and frequency-dependent strategies (i.e. the outcome of a tactic depends on the tactics of others) on the evolution of mating behaviours, usually through the use of game-theory approach (Ramsey, 2011; Smith, 1976). Alternatively, if models in evolutionary biology pay little attention to the interactions between individuals, they highlight the role of genetic architecture in determining the trajectory of evolutionary change and show that genetic variance and covariance have a potentially strong impact on evolution (Hansen, 2003; Iwasa and Pomiankowski, 1995; Iwasa, Pomiankowski, and Nee, 1991; Lande, 1976, 1981; Matessi and Di Pasquale, 1996; Walsh and Blows, 2009). Yet, mathematical resolution often requires a simplified representation of genetic architecture. For instance, in quantitative genetics, the genetic architecture of traits is described in terms of genetic variance and covariance which are supposed constant (Falconer et al., 1996, e.g. Connallon, 2015; Iwasa and Pomiankowski, 1995; Iwasa, Pomiankowski, and Nee, 1991; Lande, 1976, 1981). A set of simplifying assumptions justify this approximation (additivity, linkage equilibrium, infinite population size, multivariate Gaussian distribution of allelic effects, evolutionary equilibrium). However, in a more realistic view of the world, finite population are subject to dynamic fluctuations in the genetic variance and covariance, as a result of genetic drift and variable selection (Jones, Arnold, et al., 2003, 2007; Roff and Mousseau, 1999; Shaw et al., 1995; Steppan et al., 2002). In particular, frequency-dependent selection can increase genetic variance by favoring rare variants that differ from the most common (Sasaki and Dieckmann, 2011), and correlational selection can increase genetic covariance by favoring genetic correlation between traits (Lande, 1980; Matuszewski et al., 2014). Importantly, such changes in genetic characteristics certainly result in changes in genetic architecture and may in turn impact evolution (Debarre and Otto, 2016; Jones, Bürger, et al., 2014; Wakano and Iwasa, 2013; Wakano and Lehmann, 2014).

To better characterize the role of sexual selection in shaping mating behaviours, genetic architecture, and demography of the population, we, therefore, need to address the following questions: How do social interactions during reproduction affect the architecture of the genome? How does the evolution of genetic architecture, in turn, impact the evolution of traits and consequently affect demographic characteristics of the population? And how does the physical structure of the genome (number of genes involved, mutation rate, linkage disequilibrium) constrain this evolution?

The current challenge is thus to explicitly take into account the context-dependence effect of sexual selection (generated by mating dynamics) on the transmission of genotypes to the next generation and on the evolution of the genetic architecture.

Agent-based models (DeAngelis and Mooij, 2005) are an interesting approach to tackle this challenge, as soon as they integrate genetic transmission (Labonne and Hendry, 2010; Oddou-Muratorio and Davi, 2014; Piou and Prévost, 2012; Romero-Mujalli et al., 2019). Eco-evolutionary models using quantitative genetics showed the importance of inter individual interactions and environment on trait evolution, focusing generally on an applied or isolated question (Aguilée et al., 2013; Holt and Barfield, 2011; Labonne and Hendry, 2010; Labonne, Ravigné, et al., 2008; Oddou-Muratorio and Davi, 2014). Some genetically-explicit (i.e., allelic models) ABMs have been developed to investigate demography and evolutionary change via selection pressures from the ecological settings, but without explicitly modelling interactions between individuals, notably during reproduction (Guillaume and Rougemont, 2006; Neuenschwander et al., 2008; Peng and Kimmel, 2005). So the selective pressure on traits is *a priori* defined and therefore does not emerge from inter-individual interactions, and from trade-off between traits. By contrast, if traits values directly influence mating behaviors and survival of individuals, the selection pressure changes with the distribution of the traits values in the population; organisms thus modify their social environment and are able to respond dynamically to this change. Additionally, to our knowledge, none of these models particularly emphasizes the general role of sexual selection as a driver of traits and genetic architecture, whereas it is known to be central in evolution.

## Methods

We here give an extensive description of the model in the spirit of the ODD protocol (Grimm et al., 2006).

### An overview of the demogenetic individual based model

#### Purpose

The purpose of the present model is to investigate the co-evolution between reproductive traits under sexual selection and their genetic architecture, taking into account mating behaviour, genetic and demographic characteristics of the population. We use hereafter the term “demo-genetic agent-based model” (DG-ABM) to indicate that the present model integrates the full retroactive loops between these elements.

#### State variables and scales

This agent-based model uses a discrete temporal scale and is not spatially explicit. The time horizon of the model is in the order of tenths to a few thousand time steps; the time step can be interpreted as the minimum generation time. Three levels are considered: the gene level, the individual level, and the population level. At the gene level, genes are characterized by their alleles, which code for various genetic values depending on the number of traits simulated. They are also characterized by their position in the genome and their recombination probabilities with other genes. Such a position is invariant during the simulation because we do not wish to simulate the physical evolution of the genome. Individuals are characterized by the following state variables: identity number, sex, age, lifetime reproductive success and the genetic values of up to three traits: the gametic investment *G*, the preference *P* and the competitiveness *C*. The population level is characterized by the following state variables: population size, allelic diversity, mean allelic values and standard deviations for the various traits on each gene, mean and standard deviation of traits values and mean lifetime reproductive success. Note that the population level can be subdivided during reproduction, to create smaller mating groups and mimic spatial or temporal isolation during the breeding season.

#### Process overview and scheduling

The simulation process is described below and can be seen in Appendix A1. The model proceeds in generational time steps. Within each generation or time step, four life cycle events are processed in the following order: survival, mate choice, reproduction, mutations. Within the survival procedure, the default individual survival probability depends on population size (density dependence) and on the individual’s reproductive effort (which is the sum of the genetic values for costly traits). Within the mate choice procedure, mating groups of a user-specified size are randomly formed.

Individuals from the same mating group encounter each other (randomly, or according to their values of competitiveness), and they choose to mate or not with the encountered partner. If individuals make up a couple they will become unavailable to mate again for the present time step. Within the reproduction procedure, each parent produces gametes and the offspring are created by the random fusion of the parent gametes. Their sex is randomly assigned. Within the genetic mutation procedure, each allele might be substituted by a new one. The genetic values at each locus for each trait are drawn in independent Beta distributions by default, whose shape parameters are defined at the initialization stage (paragraph *Initialization).* This choice allows an explicit definition of the trait value, without referring to any equilibrium or average in the population. The user-defined mutation rate is assumed to be constant throughout the genome. Because survival (*S*) partly depends on reproductive effort, it can therefore also evolve, allowing individuals to potentially participate in more than one reproduction (i.e., iteroparity evolution).

### Details

#### Survival

The probability of surviving to the next reproduction event is determined as follows:

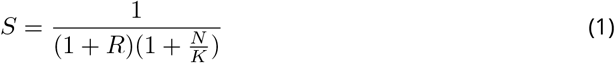

where *N* is the population size, *K* an indicator of resource limitation in the environment (akin, but not equal, to carrying capacity), *R* the sum of the genetic values of costly traits of the individual. Eq. 1 states that individual survival results from the interaction between two components. The first component is demographic 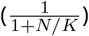, this formula corresponds to the survival rate of the population when considering that survival is solely density-dependent. The second component is 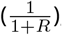, this formula states that individual survival is maximum when energy invested into reproduction is null, and that *S* decreases when R increases. Intuitively, if *N/K* = 1 (close to a demographic equilibrium) and if *R* = 1, then both components affect the survival equally. If the population size drops below *K* but *R* =1, then survival rate will increase, making R a relatively greater contributor to survival (see Fig. 1). As a consequence, the model can evolve towards semelparity (low survival, single reproduction) or iteroparity (higher survival leading to potential multiple reproductions).

**Figure 1.**
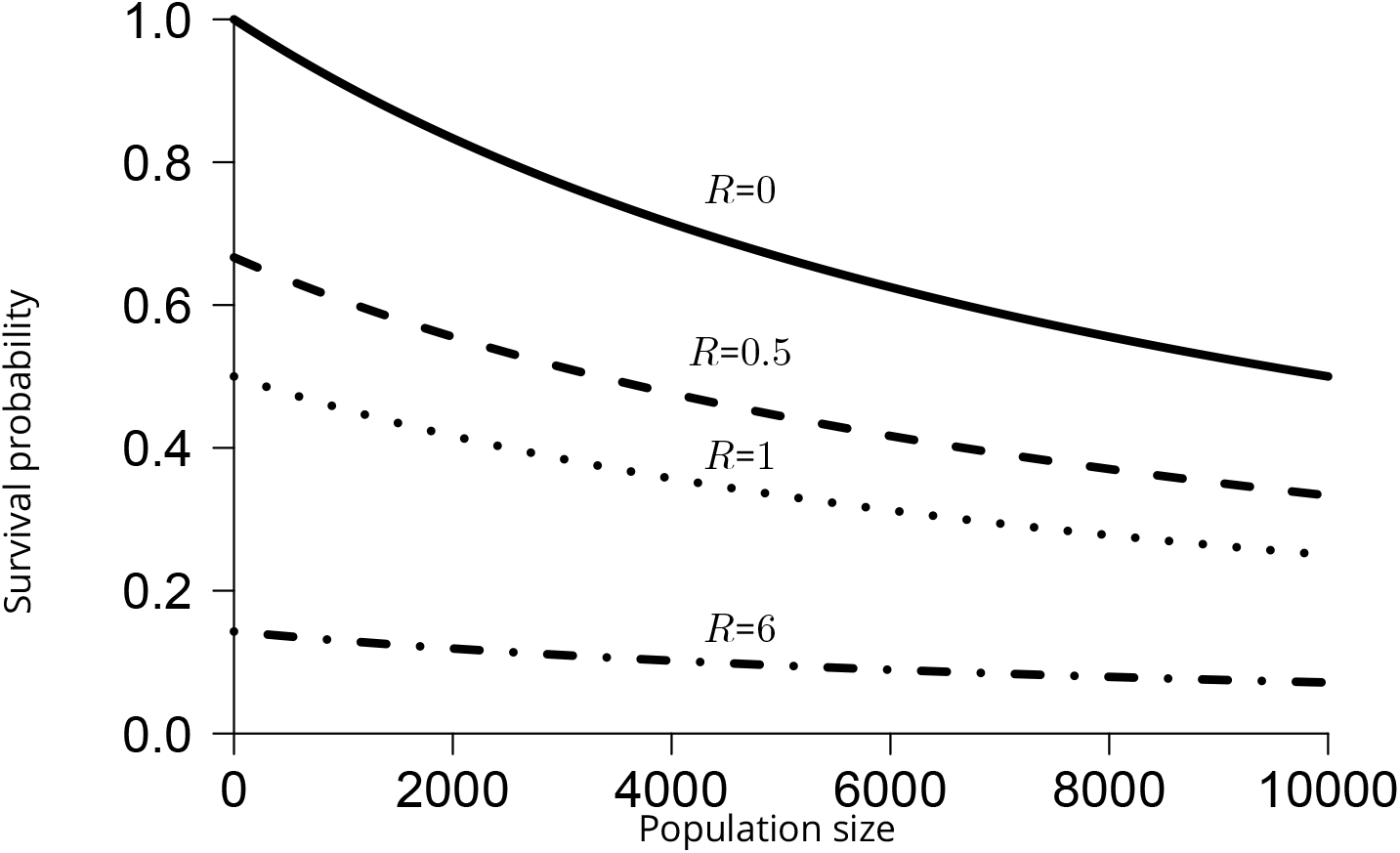
Survival probability as a function of population size for different values of *R.* with resource limitation parameter *K*=10000.

#### Mating behaviour

During reproduction, the population is first divided into mating groups of user-specified size *M*. Subdividing the population into mating groups at the time of reproduction represents the fact that individuals can potentially sample only a restricted number of mates (due to time, space, or energetic constraints, Alcock, 1991; Byers et al., 2005; Deb and Balakrishnan, 2014; Janetos, 1980; Kokko, Booksmythe, et al., 2015; Kokko and Rankin, 2006; Rintamäki et al., 1995), a central limitation when considering social interactions. Population sex ratio is 1:1 but the sex ratio within each group can vary, due to random sampling. Under mating systems involving intra-sexual competition and/or inter-sexual preference (Adler, 2011; Emlen and Oring, 1977; Kokko and Rankin, 2006), the number of potential partners encountered will be conditioned by the outcome of inter-individual interactions within the mating group, and is therefore not easily predictable. Individuals can mate only once per time step. They are thus either strictly monogamous (if they die after their first reproduction) or serially monogamous (i.e., they achieve a sequence of non-overlapping monogamous relationships and mate with a different partner at each time step). The model can represent different mating routines:

1. **random mating:** pairs of individuals from the same mating group but with different sex are formed randomly. Once a pair is formed, mating will occur. Mated individuals are not available for further mating for the current time step. Note that if *M* is not a pair number, or if all partners of one sex have mated, some individuals will remain unmated for the current time step.
2. **random encounter with preference:** pairs of individuals from the same mating group but of the opposite sex are randomly drawn, but each individual decides to accept or reject the potential partner, conditional on its own preference, and on the gametic investment of the partner. If both partners accept each other, they become unavailable for further mating for the current time step. If at least one individual rejects the mating, both of them return in the mating group and yet again two individuals are randomly drawn from the mating group. The iterative procedure stops when all individuals of one sex are depleted. To prevent the loop from spinning infinitely if individuals remain unpaired, each individual can only encounter x potential partners, x being the initial number of members of the opposite sex in the mating group. The smaller the mating group, the greater the variability of the sex ratio within the mating group, and the more variable the mating opportunities during a season (time step) will be.
3. **competitive encounter without preference:** Individuals from the same mating group but with opposite sex encounter each other in an order based on their respective values of competitiveness *C*. For each pair of individuals thus formed, mating will occur. Mated individuals are not available for further mating for the current time step. When *M* is not a pair number, or if one sex is no more available, the less competitive individuals of the mating group will remain unmated.
4. **competitive encounter with preference:** Here again, individuals from the same mating group encounter each other based on their value of competitiveness *C*. Within each pair thus formed, each individual chooses to accept the potential partner based on its preference and the partner’s gametic investment. If both partners mutually accept each other, they become unavailable for further mating for the current time step. If at least one individual rejects the mating, individuals remain free to encounter less competitive individuals.

#### Preference model

The probability that an individual will accept the mating *(Pm)* is a function of its genetic value of preference and its partner’s gametic investment (Eq. 2).

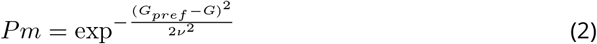

This equation indicates that individuals have a unimodal preference, i.e. they prefer a particular value of the gametic investment *(G_pref_)* with a tolerance around this value (*v*). The closer the gametic investment of the partner met is to the individual’s preferred value, the higher the probability that the individual will accept the mating (Fig. 2). The parameter *v* is set to 0.2. Other forms of preference are also available in the model and can be user specified, such as monotonously increasing preference or threshold preference.

**Figure 2.**
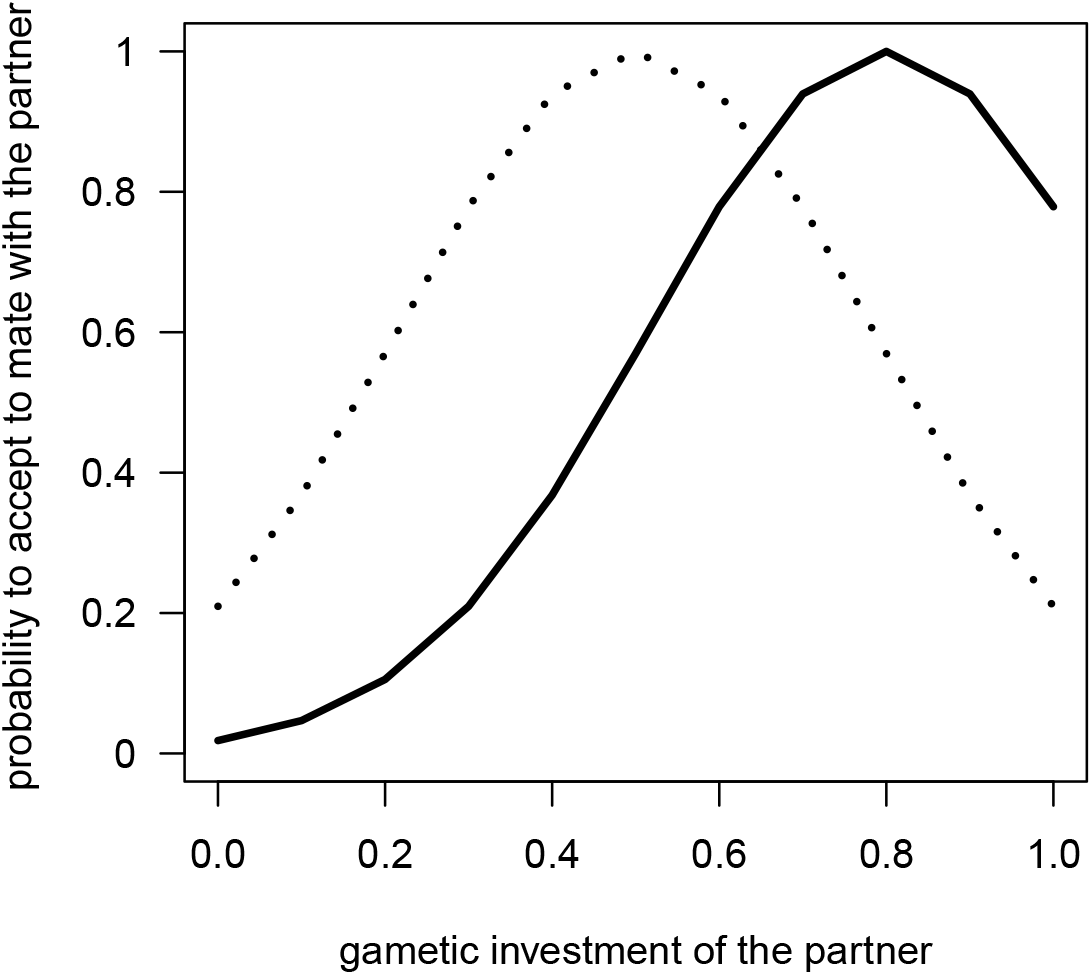
Probability that a potential partner will be accepted, as a function of the values of gametic investment of this partner for two different values of preference (*G_pref_* = 0.5 dotted line, and (*G_pref_* = 0.8 plain line). Individuals differ in the value of *G* of their most preferred mate, but all individuals share the same tolerance (*v* = 0.2).

#### Competition model

Within a mating group of size *M*, the competitiveness *C* will be used to assess a non-random order of meeting between pairs of individuals. We assume that the probability of meeting between two individuals is dependent on their respective competitiveness trait values. For instance, two individuals with high competitiveness values will probably meet first within the mating group. Then come pairs of individuals with contrasted competitiveness (one high, one low). Finally, two individuals with low competitiveness will meet at the end of the process, mostly. Pairs of individuals are thus ordered according to the product of their competitiveness. Note that if the meeting is not followed by a mating, each individual will be still available for mating. When individuals have mated once however, they are not available for further mating for the current time step and are removed from the list. Consequently, the less competitive individuals may miss reproduction if all opposite-sex partners in the mating group have already mated.

#### Reproduction

The gametes are composed of randomly chosen strands from each pair of homologous chromosomes. Recombination between successive genes may occur during the meiosis (depending on the values of recombination probabilities between genes). These probabilities are gathered in a table consisting of *n −* 1 lines, *n* being the number of genes studied. Each line *i* contains a number between 0 and 0.5 which represents the probability of recombination between the *i^th^* gene and the (*i* + 1)*^th^* gene. The number of offspring per couple is set to the average value of the parents’ gametic investments multiplied by a demographic constant. The sex of the offspring is randomly attributed.

#### Genetic mutation

Right after fecundation, mutations might occur on the offspring genome. The rate of mutation is the probability each allele has to be substituted by a new one. Alleles mutation probability is user-defined, but the default value is set to 10^-4^ mutations per loci, which is within the upper range of spontaneous mutation rates estimated at particular genes in different organisms (Haldane, 1935; Lynch, 2010; Nachman and Crowell, 2000).

#### Physical structure of the genome

All individuals share the same physical architecture of the genome, which is defined by the number of autosomes, the presence or absence of a sex chromosome, the number of genes, their location on the chromosomes and the probabilities of recombination between these genes. Each individual inherits two alleles for each gene (i.e. diploidy). Allelic effects are described by continuous values and are additive within and between loci, i.e. dominance effects are not considered, such as the value of a trait corresponds to the sum of allelic values at every locus:

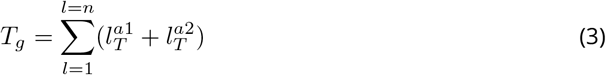

with *n* the number of loci (*n* = 100), 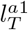 and 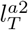 values of the first and the second allele for the trait at the loci *l*. Individual value of trait is fully determined by the genotype, meanings that environmental effects on phenotype are neglected, and heritability is thus assumed to be unity. Mutations in the model are explicitly defined as a change in allelic value, and we do not draw the effects of mutations in a distribution around a population mean value (as is done in quantitative genetics). In this framework, any mutation (i.e. any allele substitution), has an effect on the trait since the trait is built as the sum of allelic value. This approach contrasts with the other definitions of mutation (for instance, in the quantitative genetics framework), but is required to make explicit the effect of each locus on the trade-off between reproductive investment and survival that is defined at the phenotype level. We define the landscape of allelic values for preference, gametic investment, and competitiveness by Beta distributions of user-specified shape parameters. Default values are [0.65, 24.5], resulting in a right-skewed distribution such as many alleles will have small values for the traits but still, some alleles will have relatively high values for the traits (Fig. 3A). As a consequence, under this set of parameters and using 10 loci, initial trait values for gametic investment followed the distribution showed in Fig. 3B with mean 0.5 and variance 0.02. The skewed Beta distribution is selected to start from relatively low values of traits so to ensure that the initial trade-off between traits values and survival will not crash the population, and to balance the initial conditions between the investment into reproduction and the probability to survive. The user can however specify different values for the Beta distribution, and simulate uniform or bell-shaped distributions.

**Figure 3.**
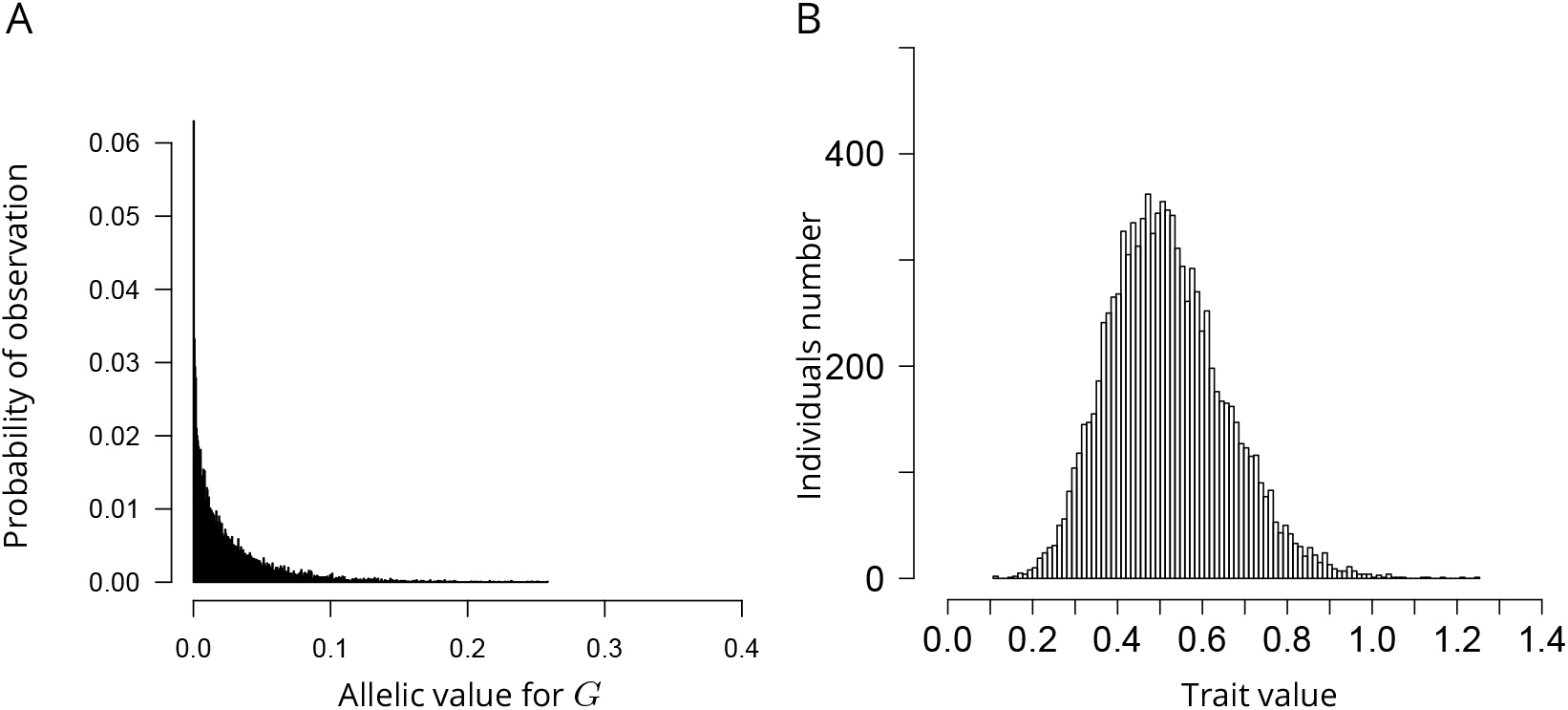
Example of allelic values distribution for the gametic investment (A) and consequent distribution of gametic investment trait values in the population (B). This distribution is obtained by drawing allelic values on the Beta distribution of shape parameters [0.65, 24.5].

#### Genetic architecture

The genetic architecture is here defined as the distribution of allelic effects along the genome, and can, therefore, vary between individuals. According to universal pleiotropy assumption (Fisher, 1930; Hill and XS Zhang, 2012; Paaby and Rockman, 2013), each allele at each locus is specified by a vector of contribution to every trait. For instance in our case, in a two traits model with gametic investment and preference, each allele has two values, one for each trait. All alleles have, therefore, a pleiotropic effect in our model. However, they can code for similar values for the simulated traits, in which case allelic correlation will be high, or they can code for contrasted values among traits, displaying low allelic correlation. This is so, because if selection acts on a trait, an allele with high correlation will not impact the fitness as will a allele with low correlation, depending on the context.

#### Initialization

To initialize a simulation, three types of parameters are specified:

- demographic parameters: the resource limitation in the environment K and the initial population size.
- mating parameters: the mating system (e.g. random mating, random encounter with preference, competitive encounter without preference or competitive encounter with preference) and the size of the mating group *M* are also required. The minimal model is run using a single trait (*G*), more complex models can be run adding mating preference (*P*) and/or competitiveness (*C*).
- parameters for the physical structure of the genome: the number of chromosomes, the number of loci in the genome, the probabilities of recombination between each contiguous pairs of loci, the maximum number of alleles per locus in the population, the distribution of allelic values, and the mutation rate.

Because initialization includes stochastic processes such as sampling in allelic effects distributions, initial states using the same parameters may vary. The model, however, allows starting several simulations out of a single initial state.

### Design concept

We here describe the general concepts underlying the design of the model.

#### Adaptation

In our model, fitness is determined by traits, social environment and demography. Individual reproductive traits can indirectly improve individual fitness: high values of *G* enhance individuals reproductive success but have a survival cost (i.e. higher probability to die per time step). High values of *C* can improve the mating success of individuals through priority access to mating partners, yet they do have a survival cost too. High values of *P* allow individuals to get a mate with high fecundity but may have an *opportunity cost* (i.e. loss of mating opportunities, De Jong and Sabelis, 1991; Dechaume-Moncharmont et al., 2016; Fawcett et al., 2012). The survival cost of *G* and *C* comes from the function of survival probability (Eq. 1). The *opportunity cost* of *P* emerges from individuals interactions. The user can however also simulates scenarios wherein *P* has a direct cost too.

#### Sensing, interactions, collectives

As previously described, during reproduction, the population is partitioned in several mating groups. Individuals sense and interact with potential sexual partners and competitors within the scale of the mating group. However, phenotypic distribution of available partners and of competitors will not affect their decision (*i.e.* no adaptive behaviour ABM wise), it only affects the outcome of interactions.

#### Evolutionary dynamics

Fitness variation drives traits evolution which in return changes the social environment (i.e. phenotypic distribution of available partners and competitors) and the demography. This feedback loop prevents ‘optimization’ in general. Rather, evolutionary dynamics can potentially converge to a pseudo equilibrium. A pseudo-equilibrium is decreed reached when evolutionary rates of traits in Haldane oscillate around zero (Hendry and Kinnison, 1999). It indicates that the product of natural selection, sexual selection, genetic drift and mutation has reached a stable balance.

#### Stochasticity

Because we use an individual-based model, most processes are inherently stochastic. Survival is, for instance, the realization of a Bernouilli random draw. Mating systems are also an important source of stochasticity. First, individuals are sampled randomly to constitute mating groups. Then, when they do not express competitiveness, individuals randomly encounter each other within each mating group. Mate choice itself is a highly stochastic process: it results from mutual acceptance of both partners, through their respective mating preference which is a probabilistic function (in most cases), so to include an error of assessment of the mating partner quality. Lastly, the transmission of genetic information is also subject to stochasticity because we represent chromosome segregation and recombination during the meiosis and we also account for mutation risk.

#### Observation

The model can either be run in graphical user interface mode or script mode. In the former, the user can select if all time steps or only a subset should be memorized. In the memorized time steps, all objects (and therefore all individuals and their genomes) are observable. A wide panel of data extractors and visualizers is then available to analyze and illustrate the simulations. In script mode, only population-level variables are recorded over time.

For each simulation, we record different type of variables to characterize mechanisms of evolution: at the population level, trait evolution is monitored by measuring the average trait values in the population and its standard deviations, and the evolutionary rate of traits in Haldanes (Hendry and Kinnison 1999, Appendix 3); genetic architecture evolution is assessed by recording the distribution of allelic values at each locus on average in the population, which gives a statistical view of the genome at the population level. From this data we calculate two indicators to characterize genetic architecture:

i. The inter-loci relative standard deviation (RSD) in genetic value indicates if all genes contribute equally or very differently to the total genetic value of traits. The inter-loci RSD is calculated as the standard deviation of genetic values between loci pondered by their mean value to look at the relative effects of mutations present in the population.
ii. The allelic correlation for two traits indicates if genes have, in average, similar effects for both traits or if, on the contrary, genes have in average different effects for each of the traits. It is calculated from the sum of the squared difference between the mean allelic values for the two traits at every locus (Appendix A2). To assess whether this allelic correlation deviates from random expectation, we used a bootstrap approach. We calculated an index corresponding to the rank of the sum of the squared difference in the distribution of the sum of the squared differences calculated by bootstrap. The bootstrap is performed *n* * (*n* — 1) times, with n the loci number. A ranking close to 0 means that the sum of differences in the average genetic values at each locus for the two traits is lower than expected by chance. A ranking close to the maximum index value means that the sum of differences in the average genetic values at each locus for the two traits is higher than expected by chance.

At the individual level: we record individuals lifetime reproductive success (fitness), individuals values of traits and individual level of allelic correlation between each couple of traits. The individual level of allelic correlation is calculated as the covariance between the allelic effects for the two traits.

#### Installation and execution procedures

The previous description of the model assumptions and mechanisms only covers a part of the settings and tools available to users in the model. A software package is available (package RUNAWAY, running under CAPSIS-4 simulation platform), allowing users to install and interact with the model, at the following address: https://doi.org/10.15454/6NFGZ9. A brief documentation, downloading and installation procedures, as well as a quick start guide can be accessed at the following address: http://capsis.cirad.fr/capsis/help_en/runaway

### Simulation experiments

To illustrate the potential of this modelling approach, we here detail two simulation experiments. Default parameters values used for all the simulations are the following: initial population size equal 10000, resource limitation is *K* = 10000, the size of the mating group *M* is set to the size of the population. The genetic map is, by default, made of 10 unlinked loci distributed over 10 chromosomes (i.e. recombination probability between adjacent locus is 0.5). The mutation rate is set to 10^-4^ mutations per loci.

### Experiment 1: coevolution between preference and gametic investment

This first example introduces the evolutionary mechanics in the model, such as the relationships between demography, individual variation, and genetics, as well as the emergent patterns in the model. We here look at the evolution of gametic investment (*G*), under two different mating systems. First, with random mating, second, with preference (*P*) driven mating, wherein individuals select partners on their gametic investment. We track how the inclusion of a non-random mating system influences the evolution of *G*, and whether *P* itself co-evolves with the former trait. We examine both the dynamic phase of the evolution during the first generations and the later convergence phase around a pseudo-equilibrium after 5000 generations, in a single population.

#### Evolution of reproductive traits

Under random mating, *G* shows a substantial evolution from its initial value (of 0.5 average in the population) to reach a pseudo equilibrium around 0.7 after 200 generations (Fig.4 A). The standard deviation around this value remains stable throughout the simulation (around 0.1). The evolutionary rate of *G* calculated in Haldanes is accordingly moderately high during the first 200 generations and then stabilize to low values. During the early phase of the simulation, we observe a clear positive correlation between individual fitness and *G*, indicating that values above the mean for this trait are beneficial (i.e. directional selection). Once at the pseudo-equilibrium, this relationship turns to a bell-shaped distribution, indicating stabilizing selection. Note that in this scenario, we also looked at the evolution of *P* as a non functional trait: this trait shows no evolution whatsoever, its average and standard deviation remaining stable throughout the simulation (Fig.4 A).

**Figure 4.**
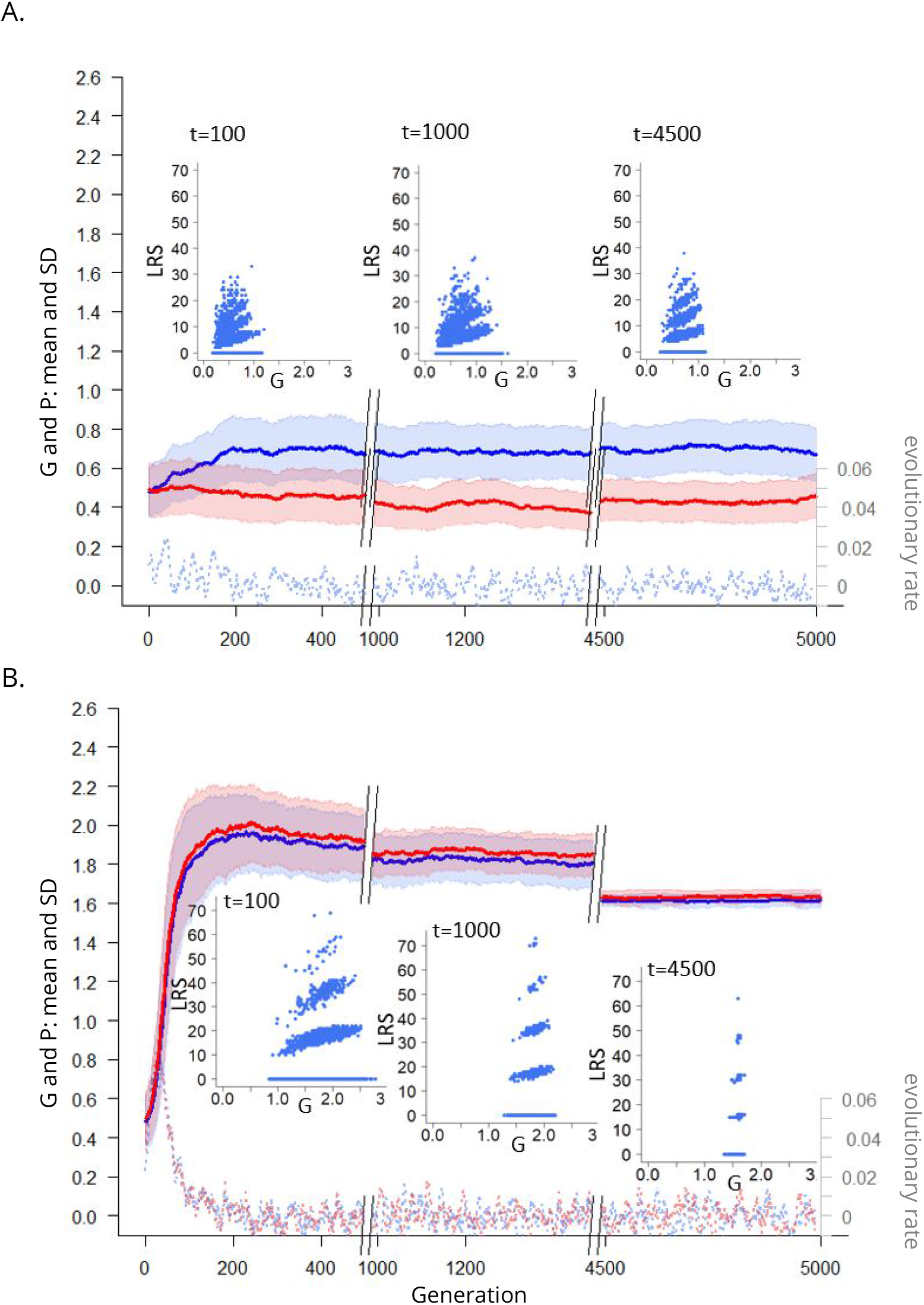
Evolution of *G* (in blue) and *P* (in red) in the population over 5000 time step, for one simulation of either (A) random mating, (B) preference driven mating. Mean values of traits are showed in thick lines and standard deviation values are represented in transparency. Also appear in the figure the evolutionary rates in Haldanes for the two traits (dashed lines) and the relationship between lifetime reproductive success (*LRS*) and *G* values at different time step during the simulation.

When mating is driven by *P*, the two traits increase quickly from their initial values (of 0.5 in average in the population) to reach maximum average values of 1.95 for *G* and 1.99 for *P* after about 200 generations (Fig.4 B). At this time, the standard deviation of the traits is maximal, around 0.2. Then the mean values of traits and their standard deviation decrease slowly overtime to reach mean values of 1.61 for *G* and 1.63 for *P* and standard deviations of 0.035 and 0.038 respectively. During the whole simulation, *P* is always greater than *G*, indicating that on average, the higher values of *G* are preferred in the population. The evolutionary rates of traits indicate a rapid evolution during the first 200 generations (with evolutionary rates that are in the upper range of values reported in the literature, Hendry and Kinnison, 1999), followed by a much slower evolution of traits. At the 100*^th^* generation, a snapshot of the fitness landscape shows that high values of *G* (of about 2) are advantageous (Fig.4 B). Later snapshots of the fitness landscapes confirm stabilizing selection, as well as an erosion of variance of *G* in the population with time (Fig.4 B). The emergence of a genetic correlation between *G* and *P* sheds some light on the rapid joint evolution of traits (Appendix A4). This correlation arises because individuals with high *P* choose individuals with high *G*, and thus, statistically, their offspring will inherit similar *P* and *G* values. It accentuates the joint evolution of the traits toward extreme values because *P* will then evolve under indirect selection (only because of the genetic correlation with *G*), while *G* is under increasing sexual selection (as *P* increases in the population) (e.g. Fisherian mechanism, Fisher, 1915; Hall et al., 2000; Lande, 1981). Interestingly, the positive correlation between the traits is transient as it reaches a maximum of 0.7 at time step 100, and then decreases to become negative (Appendix A4). Then, during the phase of stabilizing selection for the traits, the correlation oscillates around 0.

#### Genetic architecture evolution

A statistical view of the mean genome of the population shows the mean allelic effect at each locus for the two traits, in regard to the average genetic value per locus (Fig. 5). Initially, due to the sampling of allelic values in random distributions, allelic values are quasi uniformly distributed along the genome and the average allelic values per locus do not deviate from the mean (average allelic effects of loci are thus very close to 0, Fig. 5 at *t* = 1). After 5000 generations, the allelic values are not evenly distributed over the genome, some loci having strong effects; either by coding for much higher value than the average for a trait (e.g. *l*1, *l*10 for *G* and 15 for *P*) or by coding for much lower value than the average (e.g. *l*4, *l*5, *l*6, for *G* and *l*10 for *P*). The increasing variance of genetic value between loci reveals an evolution toward oligogenic architecture (and in this simulation the inter-loci RSD increases from 0.04 for *G* and 0.05 for *P* at *t* = 1 to 0.7 and 0.8 respectively). Remarkably, loci having a very high effect on one of the traits also have a very low effect on the other trait (e.g. *l*4, *l*5, *l*10). Therefore it seems that a negative correlation among allelic values for the two trait is established within loci. This observation is supported by the fact that the mean correlation among allelic values individuals’ genome is positively correlated with their lifetime reproductive success (Fig. 6). So, in the present context, alleles with opposites effects seems fixed during directional selection. This result is robust to other distribution of allelic values (uniform law, Appendix A5) but could be challenged if environmental parameters were changed (*K, M*). Within loci negative correlation may confer an adaptive advantage by allowing more combinations of traits values to be transmitted in offspring. Such modular genetic architecture could allow to better respond to the dynamic changes in demography and social environment (Clune et al., 2013; Kashtan and Alon, 2005; Lipson et al., 2002; Parter et al., 2007).

**Figure 5.**
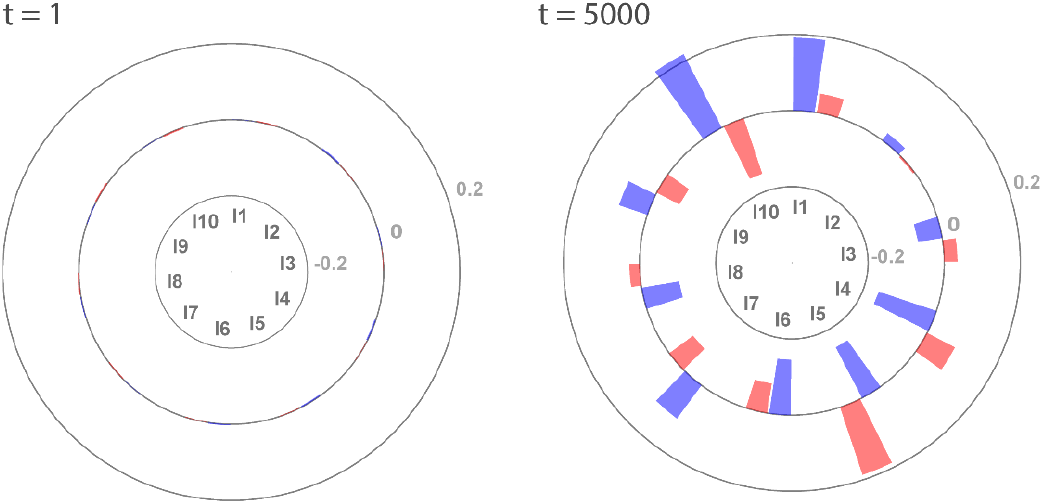
The mean allelic effects per locus in the population, at initialization (t=0) and after evolution over 5000 generation. The height of bars at each locus represents the mean value of alleles for this locus centered with respect to the average genetic value per locus for the gametic investment (blue bars) and preference (red bars). The grey lines indicate the scale.

**Figure 6.**
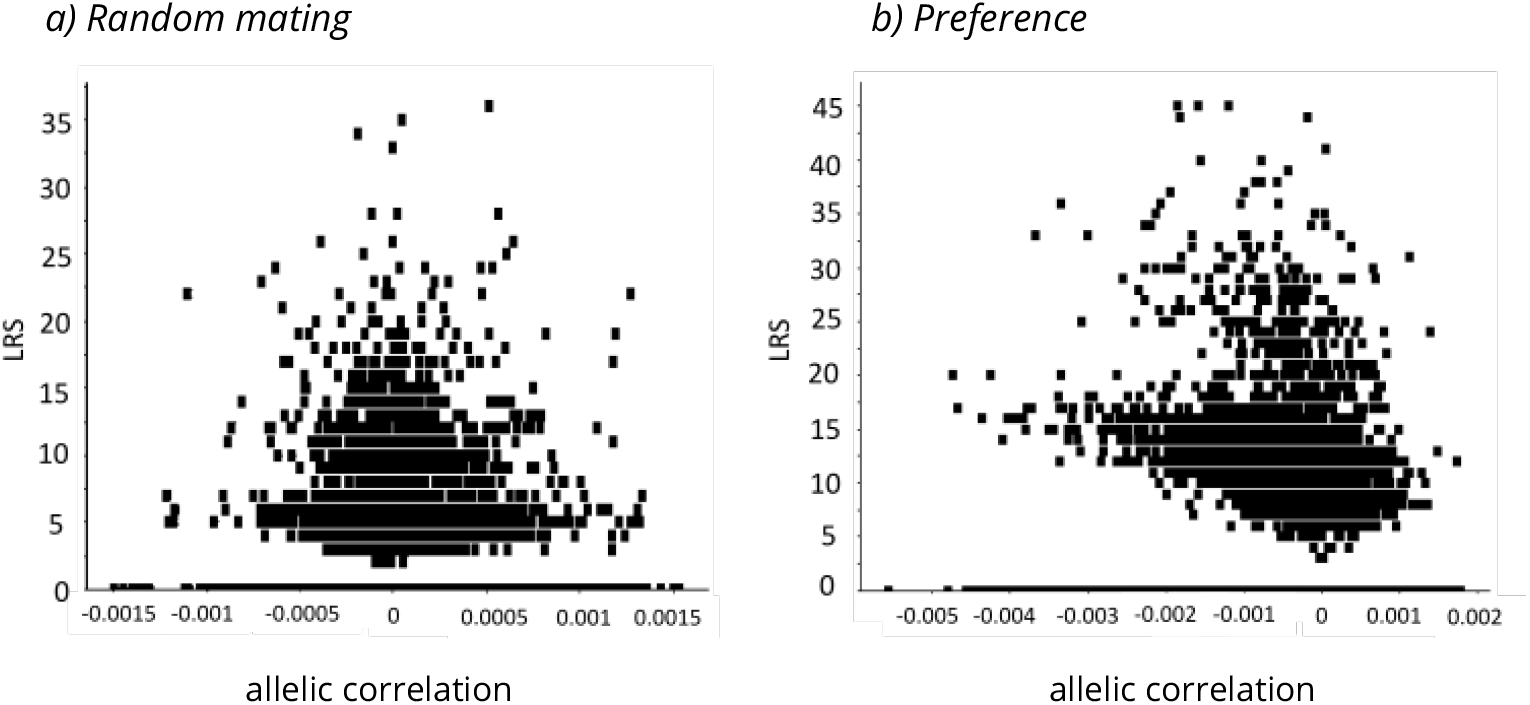
Lifetime reproductive success of individuals at timestep 50 as a function of their *G* and *P* allelic values correlation. (a) when mating is random, (b) when mating is driven by preference. In the latter case, we observe a negative correlation between individual lifetime reproductive success allelic correlation.

### Experiment 2: Effect of mating systems on traits and genetic architecture evolution

In this experiment, we generalize the above approach to 4 different mating systems, named according to the number of traits expressed in each one (random mating *G*, random encounter and preference *G* + *P*, competitive encounter without preference *G* + *C*, competitive encounter with preference *G* + *C* + *P*). Our objective is to look at the effects of mating systems on traits evolution, but also at their rippling consequences on alleles values distributions and their correlation in the genome. Here however, we perform 30 replications of simulation for each mating system, each over 5000 generations. By replicating the simulations, we investigate whether the factor we manipulate (here mating systems) leads to very homogeneous evolutionary outcomes, or whether they generate a diversified set of equilibrium. We here only focus on the final picture of the evolutionary process, without detailing the dynamics leading to it.

#### Evolution of the reproductive traits

*G* evolves toward different values depending on the scenario, which is accompanied by different population sizes at the pseudo-equilibrium (Fig. 8). When individuals express their preference (scenario *G* and *G* + *P*), *P* is favored by sexual selection, which promotes the evolution of *G* toward much higher values than in the random mating system, in which *P* evolves under genetic drift. As *G* is costly in terms of survival, its increase in the population leads to lower average survival, which translates into a lower pseudo-equilibrium population size (Fig. 8).

**Figure 7.**
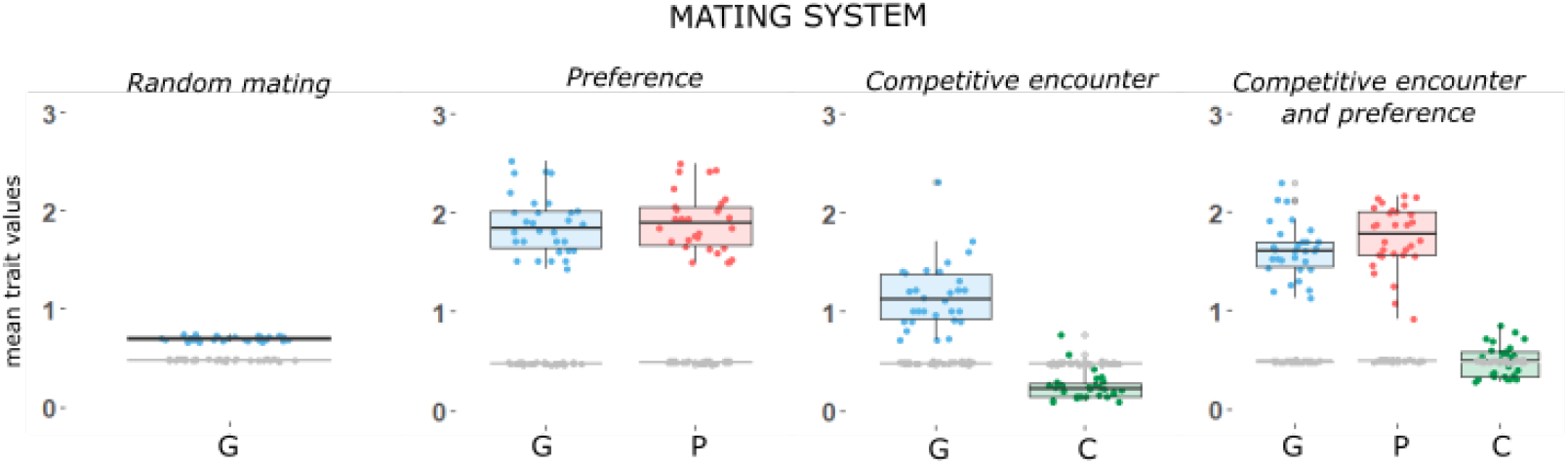
Mean values of gametic investment, preference and competitiveness, initially (in grey) after evolution over 5000 time step (in color). Four mating systems simulated: random mating (*G*), random encounter and preference (*G* + *P*), competitive encounter without preference (*G* + *C*), competitive encounter and preference (*G* + *C* + *P*). Each dot represents the main trait value in a simulation. A boxplot indicates the 2.5, 25, 50, 75 and 97.5 quantiles of the distribution among simulations.

**Figure 8.**
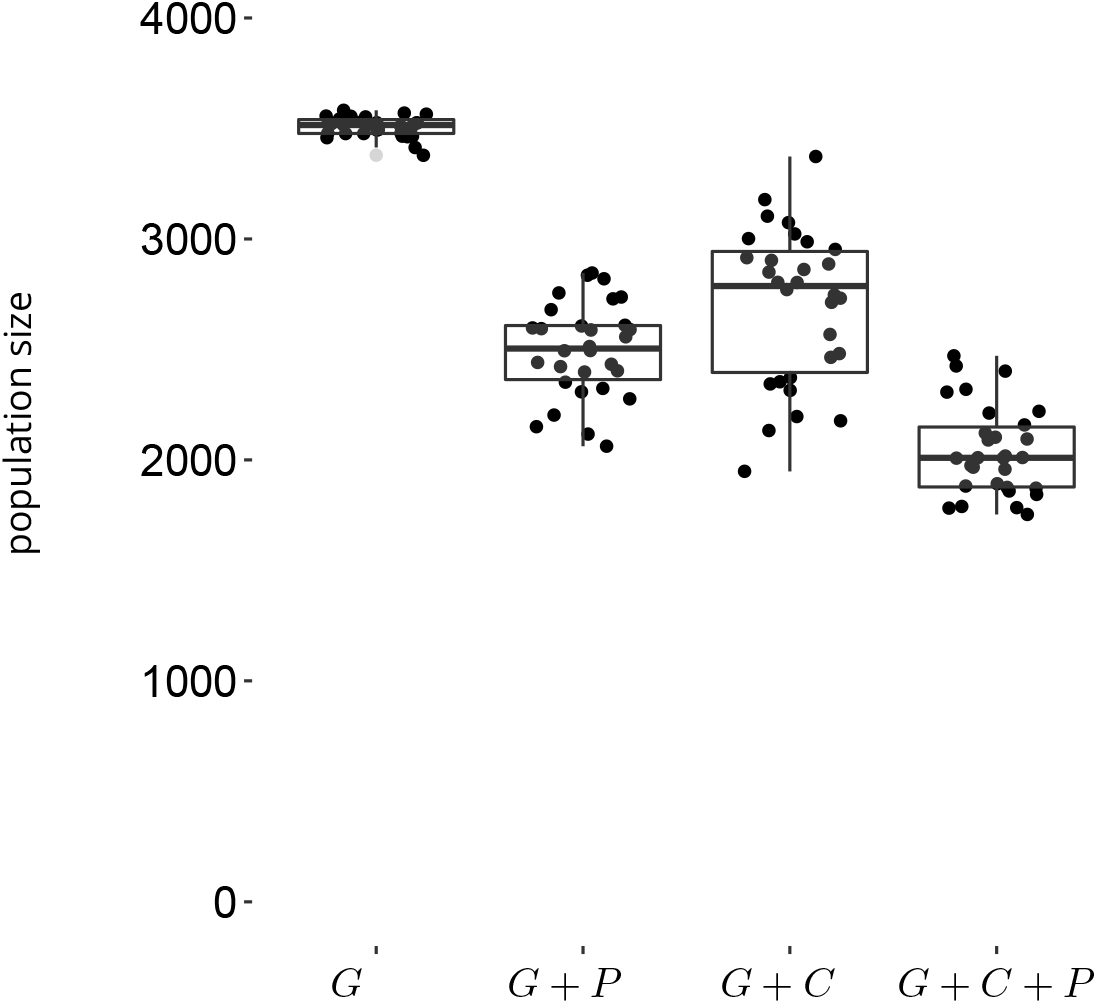
Mean population size in the different simulations after evolution over 5000 time step. Four mating systems are simulated: random mating (*G*), random encounter and preference (*G* + *P*), competitive encounter without preference (*G* + *C*), competitive encounter and preference (*G* + *C* + *P*). Boxplots indicate the 2.5, 25, 50, 75 and 97.5 quantiles of the distribution among replications.

As previously explained, the joint evolution of *P* and *G* toward extreme values is caused by the build-up of correlation between the traits (Appendix A4), which is at the core of the mechanism of Runaway selection proposed by Fisher (1915).

Competitiveness (*C*) evolution is also conditioned by the simulation scenario. *C* is selected for (despite the associated survival cost) only when individuals also express their preference *P* on gametic investment *G* (scenario *G* + *P* + *C*), because in that case, the most competitive individuals are more likely to find a suitable mate. In the scenario without preference (scenario *G* + *C*), *C* is counter selected, its cost is too great when traded off against the possible benefit of reducing opportunity costs (e.g. the risk of not finding a mate). Indeed, in the present simulations, mating group size is equal to the population size, and opportunity cost under random mating is therefore negligible.

Whereas all simulations converge to the same value of *G* at the pseudo equilibrium under random mating, as soon as there is some preference or some competitive encounter, inter-simulations variance increases. It therefore underlines the importance of the complexity of behavioural interactions on the outcome of evolution.

#### Genetic architecture evolution

Whatever the mating system investigated the relative contribution of loci to the total genetic value of each trait has somehow evolved, with some loci presenting a stronger relative contribution than in the initial situation. A difference of allelic effects distribution between mating systems is visible when looking at inter-loci relative standard deviation (RSD) of the mean allelic values (Fig. 9), the measure allows to show the difference in magnitude of genetic values between loci, independently of the differences in the values of the traits between scenarios. When mating is random, inter-loci RSD for *G* and for *P* increases, but it is slightly higher for *G*, compared to the trait *P* that is only subject to genetic drift and mutation in the random mating system (i.e., neutral trait). Under the preference driven mating system, *P* is expressed, and the inter-loci RSD of the two traits is a bit higher. When we turn to a mating system with competitive encounter without preference, inter-loci RSD further increase for both *G* and *C* traits. In contrast, when the mating system is driven by both competition and preference, inter-loci RSD for *C* is much lower, whereas the inter-loci RSD for *G* and *P* remain high (Fig. 9). These results indicate that the different mating systems, mostly due to different ways for individuals to interact for reproduction, will select for non random distribution of genetic values along the genome. This could be of course further investigated by considering different physical structure of the genome (number of chromosomes and location of genes).

**Figure 9.**
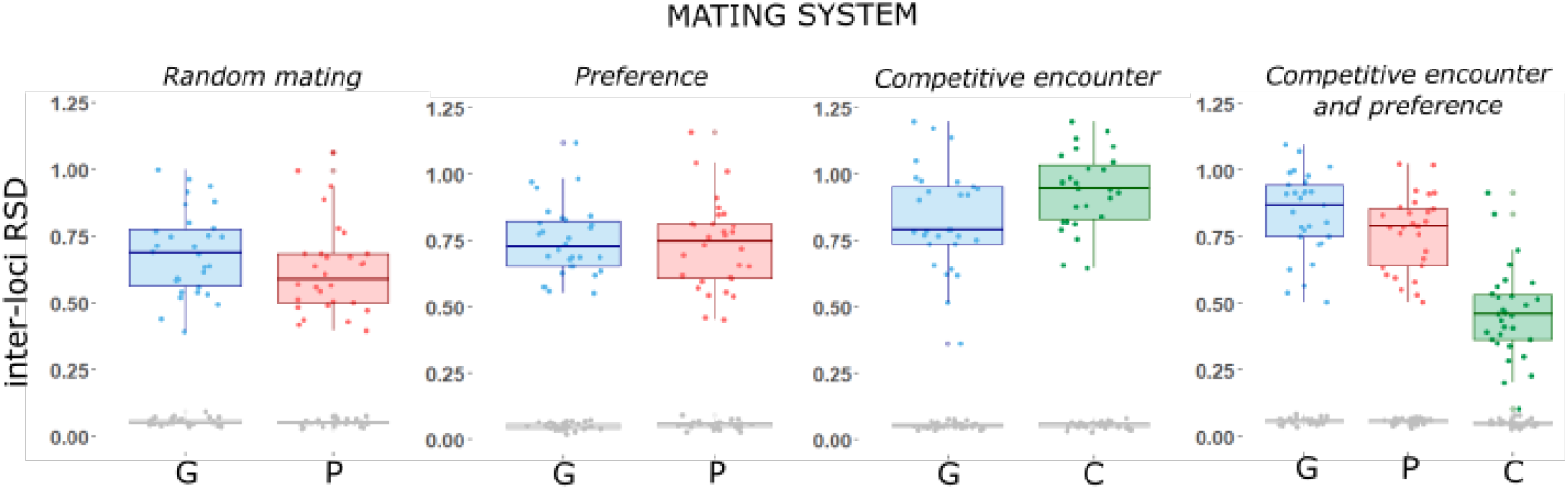
The relative standard deviation of the mean allelic values between loci for the differents traits present in the scenarios (*G*, *P*, *C*, in the first scenario the preference is not expressed), initially(in grey) and after evolution over 5000 generations (in color). The boxplots are drawn with 30 simulations of the four mating systems (random encounter, preference driven mating, competitive encounter, competitive encounter and preference driven mating) and report the mean and 95% interquartile range interval.

The pattern of correlation among allelic values within loci also evolves differently depending on the mating system (Fig. 10). When individuals mate randomly, within loci allelic correlation evolves randomly and there is no effect of initial allelic distribution: in simulations where allelic values are initially correlated by chance, the genome can evolve toward negative correlation and *vice versa,* not illustrated here.

**Figure 10.**
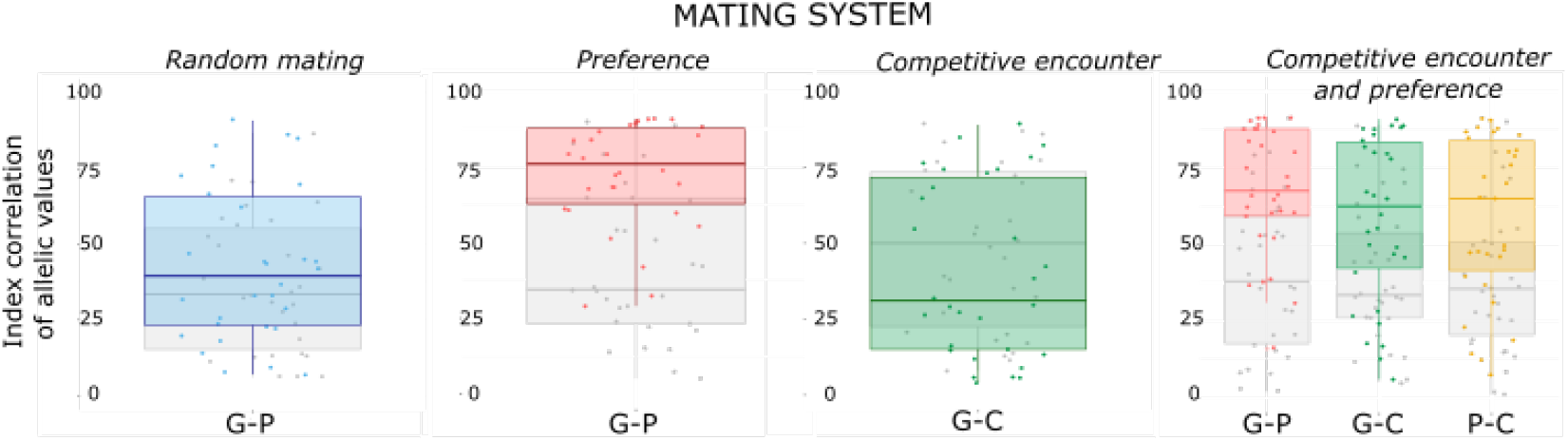
Index correlation of allelic values for the pairs of trait. The measures are showed initially (in grey) and after evolution over 5000 time step (in color). The boxplots are drawn with 30 simulations of the four mating systems (random encounter, preference driven mating, competitive encounter, competitive encounter and preference driven mating) and report the mean and 95% interquartile range interval of index values.

However, when preference matters, negative allelic correlation for *G* and *P* evolves. Remarkably, such negative correlation occurs in scenarios where there is co-evolution between the *G* and *P* and so positive global genetic correlation between the traits (scenarios *G* + *P* and *G* + *P* + *C*, Fig. 10). In contrast, when encounter is competitive but in the absence of preference, no particular organisation in the allelic effects for *G* and for *C* emerge. But here again the addition of preference select for negative correlation between allelic values and thus for a modular genetic architecture. We, therefore, observe that the mating systems mediates to some extent the evolution of the genome. Importantly, this result is robust to other mutation landscape (Appendix A5), confirming that the emergence of negative correlation between allelic values is not a computational artefact due to our draws in the distribution of allelic values.

Within the scope of this paper, we did not change the mutation rate μ, the carrying capacity *K* and the mating group size *M*. Within these settings, preference reached high values whereas competitiveness barely evolved. We suspect that it would be a whole different matter if we had modified *M*. Indeed, in situations where individuals only meet a subset of the population to find their mate (*M* < population size) the risk of not reproducing can be high, and consequently competition is expected to be stronger. In that case, traits may evolve differently; for instance, iteroparity (i.e low *G*, low *P*) can be favored as a form of “Bet Hedging” in time (Wilbur and Rudolf, 2006) and high competitiveness could be an advantage. This may, in turn, affect the evolution of genetic architecture.

## Discussion

By capturing the processes linking genes, genome, individuals, groups, population, and environment levels, DG-ABMs offer new opportunities to study evolutionary dynamics in an integrative approach. The present model is an endeavour to develop such approach in a domain where it should be especially relevant: sexual selection, an eminently context-dependent process emerging from inter-individual interactions (Kokko, Booksmythe, et al., 2015; Muniz and Machado, 2018; Otto et al., 2008). We sought, by means of examples, to illustrate how the interplay between genetics, demography, and behaviors is pivotal in predicting the evolution of traits and genetic architecture under sexual selection, whereas too often these disciplines are treated separately (Bengston et al., 2018; Rittschof and Robinson, 2014; Wilkinson et al., 2015).

Our simple simulation experiments revolved around comparing various mating systems. In these examples, wherein we kept control parameters constant along the simulations (*K*, μ, *M*, physical structure of the genome), two distinct evolutionary phases could be distinguished, and they illustrate the potential of the approach to simulate both micro-evolutionary patterns and so called evolutionary equilibrium.

The first phase is often a directional change, wherein micro-evolutionary processes can be analyzed in details, at the gene, individual or population scale. It can be perceived as a possible route towards a pseudo-equilibrium (although such pseudo-equilibrium is not certain). It depends actively on initial conditions (values of traits, based on initial genetic architecture). The direction and speed of this route are of major interest: traits evolve through optimization of fitness through selection, although we do not specify the evolutionary optimum or the adaptive landscape *per se*, contrary to most models which assume a known evolutionary optimum (e.g; Guillaume and Otto 2012; Jones, Arnold, et al. 2003; Jones, Bürger, et al. 2014; Lande and Arnold 1985; Lorch et al. 2003; Matuszewski et al. 2014; Mead and Arnold 2004). We measure the variance of fitness throughout evolution, and we can, therefore, better understand the mechanisms that lead to the selection of some trait combinations and some particular genetic architectures. In this way, the model can also be used to assess the robustness of some previous and more simple theoretical models, in a more realistic framework. For instance, we can measure the correlation between traits, relate it to fitness at each generation, and simultaneously observe whether these correlations between traits foster - or not - the building of actual non-random allelic structure within the genome. It allows us to show, among other things, that the building of genetic correlation between mating preferences and a trait - assumed by quantitative genetics approaches and at the core of Runaway selection (Fisher, 1915; Lande, 1981) - also occurs in our more complex model and explicitly arises from assortative mating process. But in our model, assortative mating is not assumed: it can potentially emerges from the evolution of preference and competitiveness, and interactions between individuals during the breeding season, achieving variable mating and reproductive success. Additionally, in the presented simulations, the rising of such genetic correlation is accompanied by the evolution of a modular genetic architecture.

In addition, the role of environmental parameters on this dynamic phase is paramount, and the model can be used to simultaneously test for the effect of habitat resource (*K*), social parameter (*M*), and genetic constraints (mutation rate μ as well as the physical structure of the genome). All of these parameters will likely affect the genetic variance, which will in turn condition the speed of evolution. The study of these effects is central to the understanding of biodiversity dynamics and resilience: for instance the model could be used to investigate variable mutation rates in time or along the genome, or to explore the effect of fluctuating resource in the environment or factors affecting the distribution of potential mating partners.

The second phase, in our current examples, is characterized by having reached an evolutionary pseudo equilibrium, the mean and variance of traits being stable within a population. Likewise, K, mutation rate, the physical structure of the genome and mating group size may have a strong impact on the outcome of the simulations. First, it can affect what are the mean values for traits, population size, and genetic architecture, but also their respective variances at equilibrium. Although these equilibria can seldom be observed in nature, where many environmental parameters fluctuate, they can nevertheless be used as a reference point to compare (qualitatively) our predictions with theoretical models. For instance, our results on genetic architecture suggest that allelic correlation can be counter selected in some situations even in the absence of functional trade-off in gene activities (Guillaume and Otto, 2012), and support the idea that selection pressures may substantially shape the distribution of pleiotropic effects among genes (Cheverud, 1996; Hansen, 2003; Pigliucci, 2008).

Additionally, predictions at equilibrium are of particular interest when we consider to what extent different replications for the same scenario may or may not converge. The difference between replications can be seen as a proxy of variance between populations, indicating that despite having the same initial conditions, environment, and constraints, two populations may diverge to some extent (when not related by dispersal, in the current model). This is of major interest for empiricists trying to investigate parallel evolution of populations experimenting similar environments (Bolnick et al., 2018; Oke et al., 2017; Schwartz and Hendry, 2007); in some cases, one should not be surprised to find substantially variable evolutionary outputs. In the core of our model, it is noteworthy to underline that such divergences mainly occur due to interactions between individuals (such as mate choice), and are therefore highly sensitive to stochastic processes. We suggest that such a mechanism could also play a major role in natural populations, stressing the importance to study and understand behavioural interactions in relation to the environment.

Arguably, the two simulation experiments only envision a small span of the possible variation in mating behaviours, the physical structure of the genome or environmental control. In its present shape, our model already includes alternative scenarios to explore, of potential interest to evolutionary and behavioural ecologists. For instance, the shape and modalities of the preference function can be easily modified by users to address fixed or probabilistic threshold responses. By modifying the physical structure of the genome, we can simulate sexual chromosomes, thereby authorizing sexual differentiation in traits genetic values. Such differences may, in turn, affect survival differently between sexes, which would, therefore, affect the operational sex ratio within mating groups, possibly modifying the strength of sexual selection, and retroactively participating in the divergent evolution of the genome between males and females (Bonduriansky, 2009; Chapman et al., 2003; Gibson et al., 2002; Hammer et al., 2008). This example indicates how all levels are naturally entangled in the individual based demogenetic approach. Notably, we have not yet included explicitly the role of parental care as well as the question of gamete size, two questions that can be central in the evolution of mating systems (Fromhage and Jennions, 2016; Kokko and Jennions, 2008; Lehtonen, Jennions, et al., 2012; Lehtonen, Parker, et al., 2016; Parker et al., 1972). For now, parental care is implicitly assumed, since the calculation of fecundity (as the mean of both partners gametic investments) somehow implies that both parents contribute to reproductive success.

The DG-ABM approach is somehow a middle ground between analytical methods and empirical approaches. On the one hand, analytical methods usually only focus on a limited sets of variables, requiring generally strong assumptions. DG-ABMs can relax some of these assumptions: for instance, there is no need to assume an equilibrium of genetic variance or stable age distribution within populations; and they are thus well suited to focus on unstable demographic or genetic situations, that are often of primary importance in evolution (Barton and B Charlesworth, 1984; Hendry and Kinnison, 1999; Kirkpatrick and Barton, 1997). On the other hand, empirical approaches can produce a wealth of patterns, on traits, on behaviour, their genetic basis, on life history trade-offs, or on population demographic and geneticstruc-tures. Only a fragment of these data are usually picked up, so to fit theoretical predictions stemming from reductive analytical models, whereas many of these patterns could help testing theoretical expectations more confidently. By allowing to generate patterns on several levels, we hope that DG-ABM way can help strengthen the links between theory and observations. Obviously, a central concern in this matter lies in the validity of the mechanisms and assumptions we made, and how well they allow to recreate either theoretical facts or natural patterns. Such question cannot be addressed independently from the simulations scenarios built by the user, since the sets of parameters (physical structure of the genome, ecological parameters, type of mating system, etc.) will directly impact the results. We can however give examples on how to connect some outcomes of the model to either theoretical or empirical expectations. For instance, when we investigated the coevolution between sexual preference and gametic investment, both traits evolved rapidly at the beginning - since we were unlikely to start from an equilibrium. But during this phase of evolution, genetic variance increased. It later decreased when the population reached a pseudo-equilibrium. We thus reproduced a known result, which is the transient runaway as trait and preference eventually goes to a new steady state (De Jong and Sabelis, 1991). Additionally, the pseudo-equilibrium reached was admittedly stable. Lande (1981) predicted in his model that stability in this case was only ensured when the ratio of trait-preference covariance over trait variance in the population was equal or below 1 (in case of equal mutation variance input for both traits). Above that value, the equilibrium becomes unstable. The said ratio was observed to reach values up to 0.8 over our simulations, but it never exceeded 1. We also explored different physical structure of the genome (with different loci numbers, linkage disequilibrium, mutation rates; data not shown) but in the range of parameters explored, no simulation produces runaway selection toward unstable equilibrium. The non-occurrence of unstable equilibrium could be due to finite population size and the resulting genetic drift, potentially erasing the genetic correlation (Nichols and Butlin, 1989); or it could be caused by the opportunity cost that emerges from the individual interactions in the model and slows down the evolution of preference (Kokko, Booksmythe, et al., 2015).

On a more empirical perspective, on the same example, we were able to measure evolutionary rates of about 0.06 Haldanes over 10 generations during the phase of rapid evolution with preference, and then of 0.005 (over 10 generations) at the pseudo equilibrium. These values are in agreement with empirical studies: for instance, Karim et al. (2007) reported rates of male color change following an experimental introduction that ranged from 0.01 to 0.031 Haldanes over 13–26 generations. These values are close to those we observe during the dynamic phase of evolution.

## Conclusion

Beyond the question of co-evolution between gametic investment and preference, the DG-ABM modelling approach allows to make much more comprehensive predictions on biological systems: for instance, instead of solely focusing on the evolutionary equilibria of trait values, we here provide a complete picture of what such equilibria entail also in term of genetic architecture and demography. Namely, for such dynamics to occur, we demonstrate that demographic characteristics will also reach non-random values (level of iteroparity and survival, effective numbers of breeders, population size) and that this may come with the building of non-random allelic structure within the genome. Such projection also allows capturing more efficiently the evolutionary constraints that will control the outcome of adaptive evolution (genetic architecture, phenotypic correlation, demographic processes). We hope that the present contribution can motivate further work on the link between the physical structure of the genome and variations in mating systems or life histories (e.g. D Charlesworth and Wright 2001; Lamichhaney et al. 2016; Misevic et al. 2006; Plomion et al. 2018; Sinervo and Svensson 2002).

## Acknowledgements

The present model was coded on the CAPSIS-4 modelling platform (http://capsis.cirad.fr, Dufour-Kowalski et al. 2012) maintained by F. de Coligny and N. Beudez, who also helped in developing parts of the code. We also thank A. Courtiol for thoughtful suggestions on an earlier version of the model. Version 4 of this preprint has been peer-reviewed and recommended by Peer Community In Evolutionary Biology (https://doi.org/10.24072/pci.evolbiol.100112).

## Data accessibility

Simulation results and R scripts are available on demand. A software package is available (package RUNAWAY, running under CAPSIS-4 simulation platform), allowing users to install and interact with the model, at the following address: https://doi.org/10.15454/6NFGZ9. A brief documentation, downloading and installation procedures, as well as a quick start guide can be accessed at the following address: http://capsis.cirad.fr/capsis/help_en/runaway

## Appendix

### A1. The model

**Figure A1.**
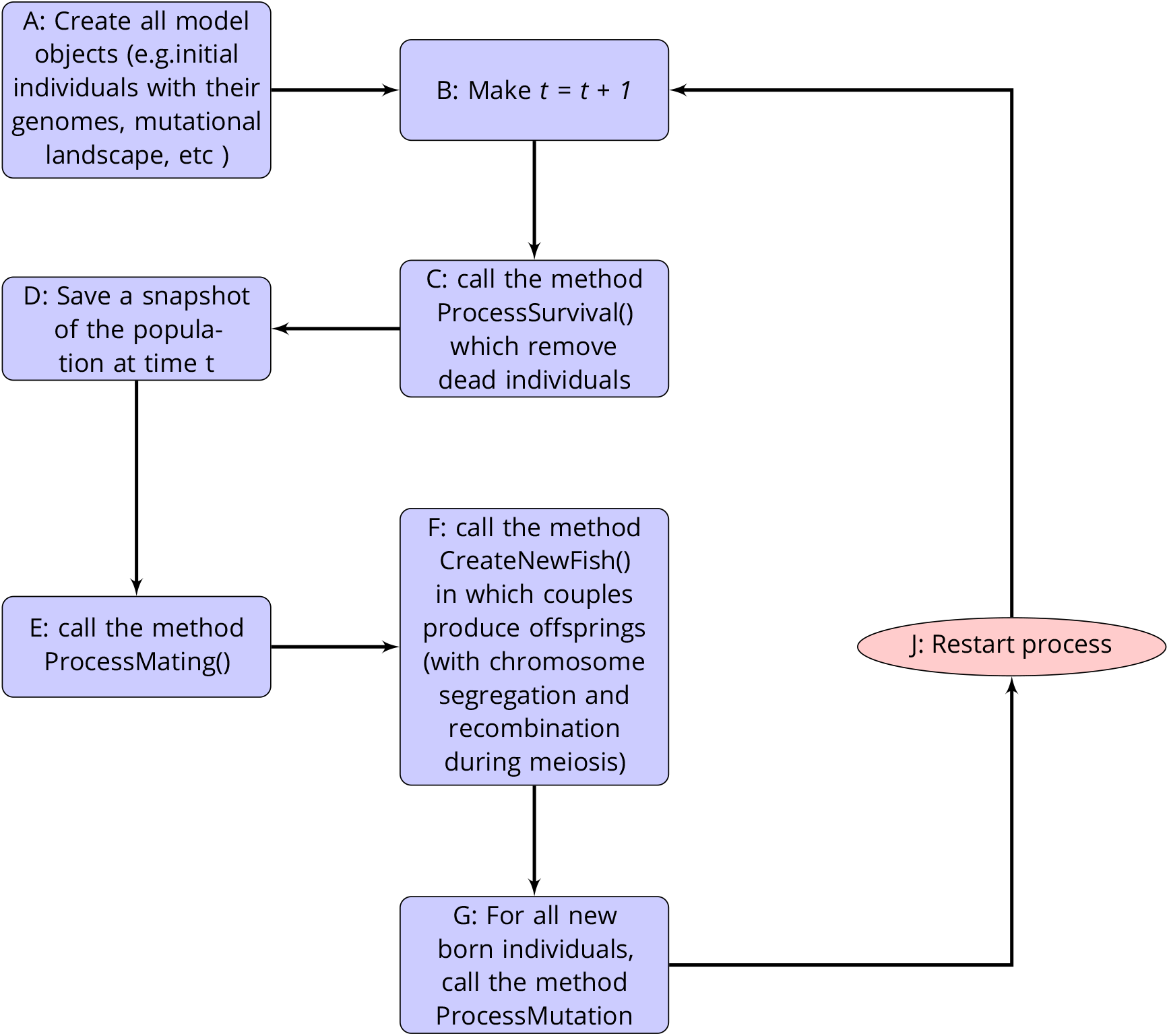
Flowchart showing the order of the processes implemented in the model.

### A2. calculation of the allelic correlation index by bootstrap

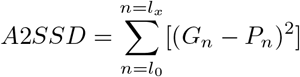

with *G_n_* and *P_n_* the mean allelic values at loci *n* for *G* and *P* and *x* the number of loci. SSD is the sum of square differences between allelic values for the traits at every loci. The pleiotropy index corresponds to the rank of the sum of the squared difference in the distribution of the sum of the squared differences calculated by bootstrap.

### A3. Evolutionary rate in Haldanes

We measure evolutionary rate in Haldanes using the same formula as Hendry and Kinninson (1999):

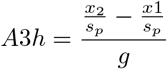

where *x_2_* and *x*_1_ represent mean trait values for the single population at two different times, *s_p_* is the pooled standard deviation 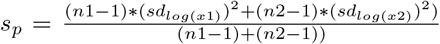, and *g* is the number of generations between the two different times.

### A4: supplementary figures

**Figure A4.**
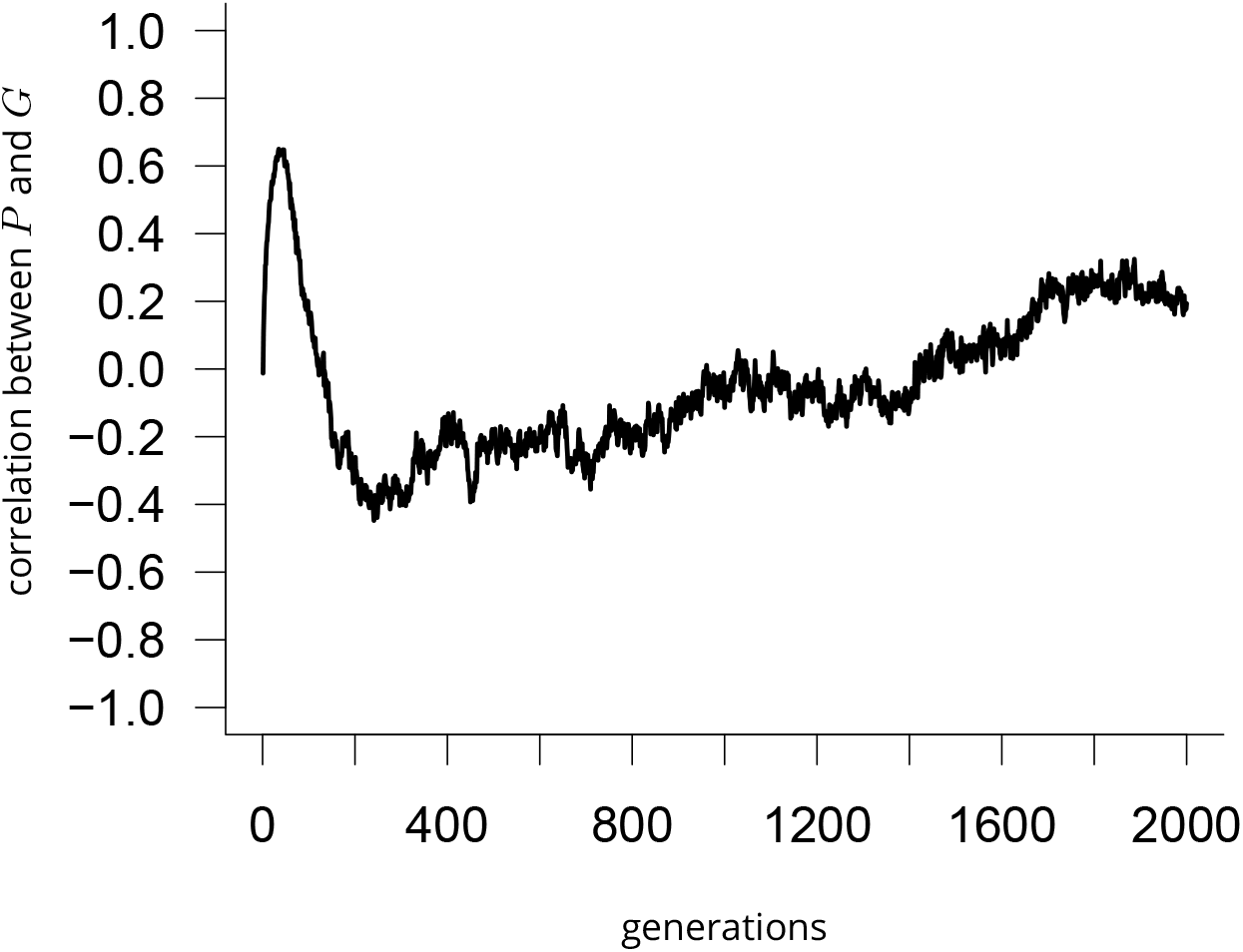
Individual genetic correlation between *G* and *P* values in the population, for one simulation of the preference driven mating system (*G* + *P*).

**Figure A5.**
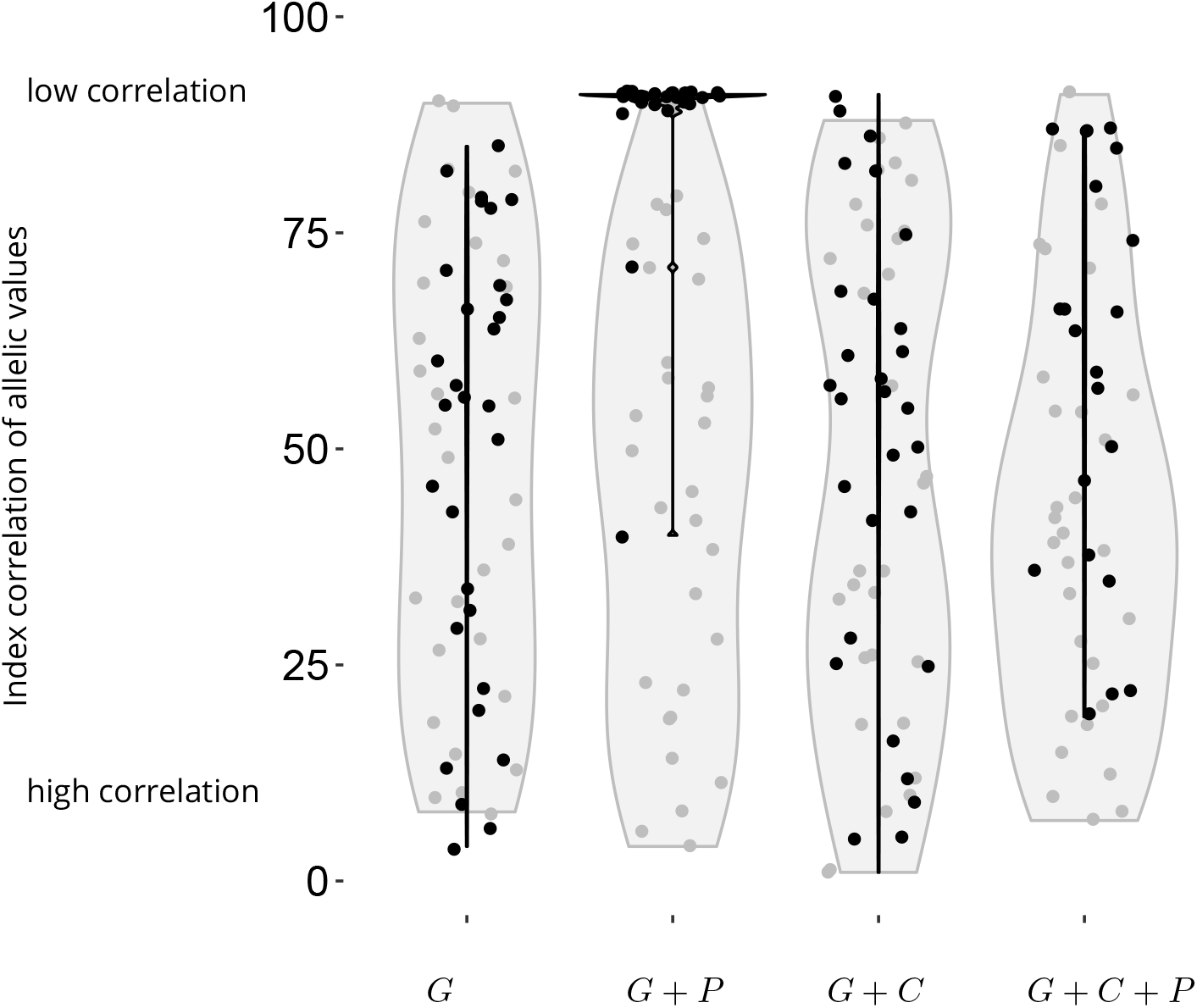
Index correlation of allelic values for *G* and *P* with an uniform distribution of allelic values. The measures are showed initially (in grey) and after evolution over 5000 time step (in black) for 30 replications for each mating system (random encounter without preference, random encounter and preference, competitive encounter without preference, competitive encounter and preference). Greyed areas indicate density probability for the distribution of points in the variation range. The mutational landscape follows an uniform distribution.

